# TORC2-regulated sterol redistribution mediates recovery from membrane perturbation by small amphipathic molecules

**DOI:** 10.1101/2024.10.18.618785

**Authors:** Maria G. Tettamanti, Paulina Nowak, Beata Kusmider, Jennifer M. Kefauver, Vincent Mercier, Aurélien Roux, Robbie Loewith

## Abstract

To maintain plasma membrane (PM) integrity, cells need to acutely regulate PM lipid composition. The Target Of Rapamycin (TOR) complex 2 is a protein kinase that acts as a central regulator of PM homeostasis, but the mechanisms by which it monitors and reacts to membrane stresses are poorly understood. To address this knowledge gap, we characterized a family of amphiphilic molecules that physically perturb PM organization and in doing so inhibit TORC2 in yeast and mammalian cells. Using fluorescent lipid associated reporters in budding yeast, we show that these small molecules first cause a transient increase in the amount of biochemically accessible ergosterol at the PM. Contemporaneous TORC2 inhibition stimulates a rapid removal of accessible ergosterol from the PM by the PM-ER sterol transporters Lam2 and Lam4, necessary for TORC2 reactivation. Thus, we show that TORC2 acts in a feedback loop to control active sterol levels at the PM and introduce sterols as possible TORC2 signalling modulators.

## Introduction

The plasma membrane (PM) is an interface between the cell and its surroundings. It serves both as a selectively permeable barrier and as a signalling platform (Leonard, Loose and Martens, 2023). The involvement of the PM in a large collection of cellular processes is enabled by its lateral compartmentalization into functional domains (Honigmann and Pralle, 2016; Lu and Fairn, 2018). This is dependent on the biophysical properties of the PM, and thus on its complex protein and lipid composition and configuration. To maintain optimal PM properties, cells actively regulate their lipid repertoire. (van Meer, Voelker and Feigenson, 2008; Harayama and Riezman, 2018) Sterols, such as ergosterol in yeast, are a good example of such regulation. Sterols are major modulators of physical membrane properties, such as rigidity and phase behaviour, and they are especially enriched at the PM (Dufourc, 2008; Doole *et al*., 2022; Varma and Deserno, 2022). As important factors in membrane microdomain organization(Kefauver *et al*., 2024), most sterol molecules in membranes are cloistered by phospho- and sphingolipids, and only the excess fraction is free and accessible for biochemical processes (Radhakrishnan and McConnell, 2000; Ohvo-Rekilä *et al*., 2002; Sokolov and Radhakrishnan, 2010; Lange *et al*., 2013; Das *et al*., 2014; Lange and Steck, 2020). Sterol levels in internal membranes vary and require tight regulation (van Meer, Voelker and Feigenson, 2008). The redistribution of sterols between cellular membranes depends on various vesicular and non-vesicular transport routes, of which cytosolic and membrane-contact-sites lipid transport proteins are the most efficient (Kaplan and Simoni, 1985; Baumann *et al*., 2005; Dittman and Menon, 2017; Wong, Gatta and Levine, 2019; Steck, Tabei and Lange, 2021).

Target Of Rapamycin Complex 2 (TORC2) is one of two functionally and structurally conserved protein complexes containing the essential protein kinase TOR (Heitman, Movva and Hall, 1991; Loewith *et al*., 2002; Wullschleger, Loewith and Hall, 2006). While specific inhibition by rapamycin has facilitated characterization of TORC1, TORC2 is rapamycin-insensitive and thus its characterization has relatively lagged (Loewith *et al*., 2002; Gaubitz *et al*., 2015). Recent studies have positioned TORC2 as a central regulator of PM homeostasis which reacts to changes in the biophysical properties of PM: the application of various orthogonal stimuli that cause an increase in membrane tension and/or other stress - i.e. hypo-osmotic shock, sphingolipid biosynthesis inhibition - also cause an increase in TORC2 activity, while those that reduce membrane tension - hyper-osmotic shock and treatment with palmitoylcarnitine (PalmC) - rapidly inactivate TORC2 (Berchtold *et al*., 2012; Riggi *et al*., 2018). TORC2 regulates several processes that affect turgor pressure and/or plasma membrane properties, via its primary effector, the AGC family kinase Ypk1, (reviewed in (Roelants *et al*., 2017; Thorner, 2022)). The retrograde PM-ER sterol transporters Lam2 and Lam4 are amongst the downstream effectors regulated by Ypk1 (Roelants *et al*., 2018; Topolska *et al*., 2020), reinforcing TORC2’s role in PM biophysical homeostasis. Consistent with this role, acute chemical inhibition of TORC2 increases PM tension, while Ypk1 hyperactivation decreases it (Riggi *et al*., 2018). Collectively, these data have recently led us to propose that TORC2 functions in a mechanosensitive feedback loop to maintain the biophysical homeostasis of the PM (Riggi, Kusmider and Loewith, 2020). This role of TORC2 is likely conserved in higher eukaryotes, as evidenced by reports showing that the major mTORC2 substrate Akt is regulated downstream of changes in membrane tension and that mTORC2 signalling feeds back to control PM properties (Kippenberger *et al*., 2005; Diz-Muñoz *et al*., 2016; Roffay *et al*., 2021; Ono, Matsuzawa and Ikenouchi, 2022).

How TORC2 monitors PM status and adapts signaling to react to different stresses remains poorly understood. In budding yeast, TORC2 stimulation by PM stress is dependent on the tension dependent release of the regulatory proteins Slm1 and Slm2 from the eisosomal compartment, a stress sensitive PM domain scaffolded by BAR domain proteins (Berchtold *et al*., 2012; Appadurai *et al*., 2020; Lanze *et al*., 2020; Sakata *et al*., 2022; Kefauver *et al*., 2024). Counterintuitively, TORC2 inhibition seems to be independent of Slm1/2 relocation and instead correlates with the formation of PI(4,5)P2-containing PM invaginations (Riggi *et al*., 2018). In neither case are the molecular mechanisms of TORC2 regulation well understood. During hyperosmotic shock, transient TORC2 inhibition is paired with a transient activation of the Hog1 MAPK cascade. These changes in TORC2 and Hog1 activities converge on the production and efflux of the osmolyte glycerol to mitigate the offset of cell volume and turgor and ultimately restore homeostasis (Lee *et al*., 2012; Muir *et al*., 2015; Riggi *et al*., 2019; de Nadal and Posas, 2022). We previously identified the small amphiphile palmitoylcarnitine (PalmC) as an indirect TORC2 inhibitor (Riggi *et al*., 2018). PalmC inserts into the PM, and like hyperosmotic shock, reduces PM tension and transiently inhibits TORC2, but, surprisingly, does not activate Hog1 (Riggi *et al*., 2018). This suggests that PalmC triggers a PM stress that is mechanistically distinct from hyperosmotic shock stress, and likely requires different adaptive mechanisms to resolve. Building on this observation, we sought to use PalmC as a tool to better understand the physicochemical properties of the PM that are sensed upstream of TORC2. We found that exposure to small amphipaths causes a transient increase of free sterol molecules at the PM, which correlates with TORC2 inhibition. This in turn stimulates a rapid removal of free PM ergosterol, dependent on the START-domain proteins Lam2 and Lam4 (Gatta *et al*., 2015; Murley *et al*., 2017; Topolska *et al*., 2020), which is necessary for TORC2 reactivation. Thus, our data describe a feedback loop between TORC2 activity and free sterol levels at the PM, highlighting TORC2’s role in regulating PM biophysical properties and domain organization.

## Results

### PalmC partitions into the PM to inhibit TORC2

We have previously reported that the small amphipathic molecule PalmC (Fig. 1A, grey panel) acts as an indirect inhibitor of TORC2 by directly interacting with the PM. This correlates with the formation of PI(4,5)P_2_ -containing PM invaginations and a decrease in membrane tension.

**Fig. 1:**
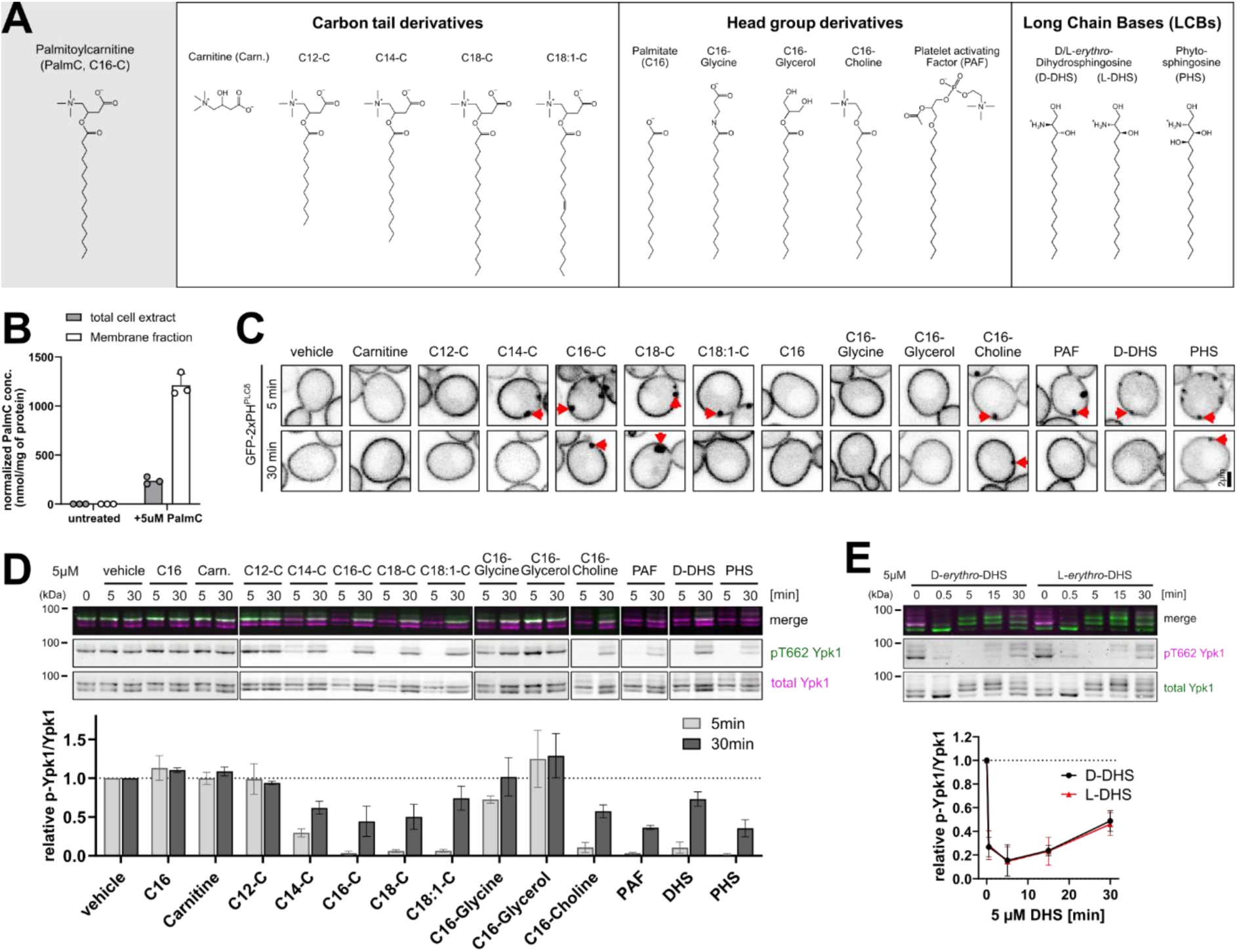
A family of small amphipaths targets the plasma membrane to inhibit TORC2. ***A)*** Structure formulas of tested PalmC derivatives. ***B)*** Mass spectrometry analysis of whole cell and membrane fraction metabolite extracts from untreated yeast cells or yeast cells treated with 5 µM PalmC. The amount of PalmC was normalized to the protein content of each extract. Mean and SD were plotted. ***C)*** Live cell fluorescence microscopy of WT cells expressing GFP-2xPH^PLCδ^ from plasmid. Cells were treated with the indicated substances at 5 µM for 5 or 30 min. Red arrows indicate PM invaginations. ***D,E)*** Western blot analysis of TORC2 activity in WT yeast cells. Cells were treated with ***D)*** 5 µM different PalmC derivatives, or ***E)*** with 5 µM D-*erythro-* or L-*erythro-*DHS for the indicated amounts of time, and TORC2 activity was assessed by relative phosphorylation of Ypk1. Mean and SD.

Mass spectrometry analysis of extracts from yeast cells treated with PalmC (in which PalmC is not a naturally occurring metabolite) showed that PalmC accumulates in the membrane fraction (Fig. 1B). Indeed, previous *in vitro* studies have revealed that PalmC virtually completely partitions into lipid bilayers in aqueous solution (Requero, Goni and Alonso, 1995; Goñi, Requero and Alonso, 1996). We observed that the kinetics and magnitude of PalmC effect correlated closely with the absolute amount rather than the concentration of PalmC administered to a population of cells, and thus with culture density (Fig. S1A). This difference cannot be attributed to differences in culture stage, or secreted factors, since it is observed in cells from the same logarithmically growing culture, which were pelleted and resuspended to different culture densities (OD_600nm_) in fresh media. Thus, controlling culture density offers a way to fine-tune PalmC effect at sub-CMC levels. We conclude that the effect of PalmC must be a consequence of its propensity to partition directly into cell membranes, in our case most probably the plasma membrane.

### A family of small amphipaths with similar properties recapitulates the effect of PalmC

To characterize which molecular features of PalmC are necessary for its effect on the PM and TORC2, we performed a Structure Activity Relationship (SAR) analysis. We screened a panel of derivatives, differing from PalmC in either the length (or saturation) of their fatty acyl chain, or in their headgroup moiety (Fig. 1A), for their ability to recapitulate the PalmC effect.

First, we tested if any of these derivatives were able to cause PM invaginations both in live yeast cells expressing the PM PI(4,5)P_2_ FLARE (fluorescent lipid-associated reporter) GFP-2xPH^PLCδ^ (Kavran *et al*., 1998) (Fig. 1C), and in a high throughput microscopy screen with fixed cells expressing mCherry-2xPH^PLCδ^ (Fig. S1B). Both approaches showed that a carbon tail of at least C14 was necessary to induce PM invaginations, and the effect was more substantial and lasted longer in carnitine derivatives with a carbon tail length of C16 or greater. In general, a cationic (choline, LCBs) or zwitterionic (carnitine, PAF (as previously described (Kennedy *et al*., 2014)) headgroup was necessary for induction of PM invaginations, while amphipaths with uncharged headgroups had no effect.

Next, we checked the effect of our derivatives on TORC2 activity. Substances that caused PM invaginations also inhibited TORC2-dependent phosphorylation of Ypk1 (Fig. 1D) (but not TORC1 phosphorylation of Sch9 (Fig. S1C)). The inhibitory effect correlated with the effect size and duration of PM invaginations (Figs. 1C, S1B). The inhibition of TORC2 by small amphipaths is conserved in mammalian cells, where PalmC (in a dose dependent fashion) and derivatives that were effective in yeast trigger the loss of mTORC2 mediated phosphorylation of Akt (Fig. S1D,E). We continued to characterize the effects of small amphipaths in budding yeast.

### The effects of small amphipaths are independent of metabolization

In our SAR screen, we had also included the long chain sphingoid bases (LCBs) dihydrosphingosine (DHS) and phytosphingosine (PHS), which resemble PalmC in structure and effect, but unlike PalmC are metabolites naturally found in yeast. It had been reported previously that the addition of PHS leads to TORC2 inhibition (Lucena *et al*., 2018); however, in this previous work, it was postulated that processing of long chain base into more complex sphingolipid species was required for TORC2 inhibition, a model at odds with our proposition that these amphiphiles act directly on the PM to trigger TORC2 inhibition. To challenge these models, we exploited a non-physiological / non-metabolizable enantiomer of DHS – L-*erythro-*DHS (Watanabe *et al*., 2002) (Figure 1A, right panel). The addition of L-*erythro-*DHS triggered rapid (after 30 seconds) TORC2 inhibition with kinetics that were indistinguishable from D-*erythro-*DHS (Fig. 1E), and also induced PM invaginations (Fig. S1B). Thus, LCBs (and likely other amphipaths) directly act on the PM and inhibit TORC2 without the need to be metabolized. For the remainder of the study, PalmC was used as a PM targeting drug, since it gave us the most robust effect in the SAR screen (see Fig. 1C,D, S1B).

### Small amphipaths cause selective redistribution of PM ergosterol

Perturbation by PalmC results in PM remodelling, with PI(4,5)P_2_ accumulating transiently in giant PM invaginations (Riggi *et al*., 2018). To characterize better how PalmC affects the distribution of different lipid species, we complemented our assessments of PI(4,5)P_2_ behaviour (mCherry- or GFP-2xPH^PLCδ^), by additionally observing phosphatidylserine (PS; GFP–C2^Lact^ (Yeung *et al*., 2008)), PI(4)P (mCherry-P4C (Luo *et al*., 2015)), and ergosterol (yeGFP-D4H (Maekawa, Yang and Fairn, 2016; Marek, Vincenzetti and Martin, 2020)). We observe that in addition to PI(4,5)P_2_, both PS and PI(4)P also localize to the same PM invaginations after PalmC addition (Fig. 2A, S2A, left and middle panels), where they accumulate to a similar degree as PI(4,5)P_2_ (Fig. S2B). There is no decrease in colocalization with PI(4,5)P_2_, neither for PS nor for PI(4)P (Fig. 2A, left and middle plots).

**Fig. 2:**
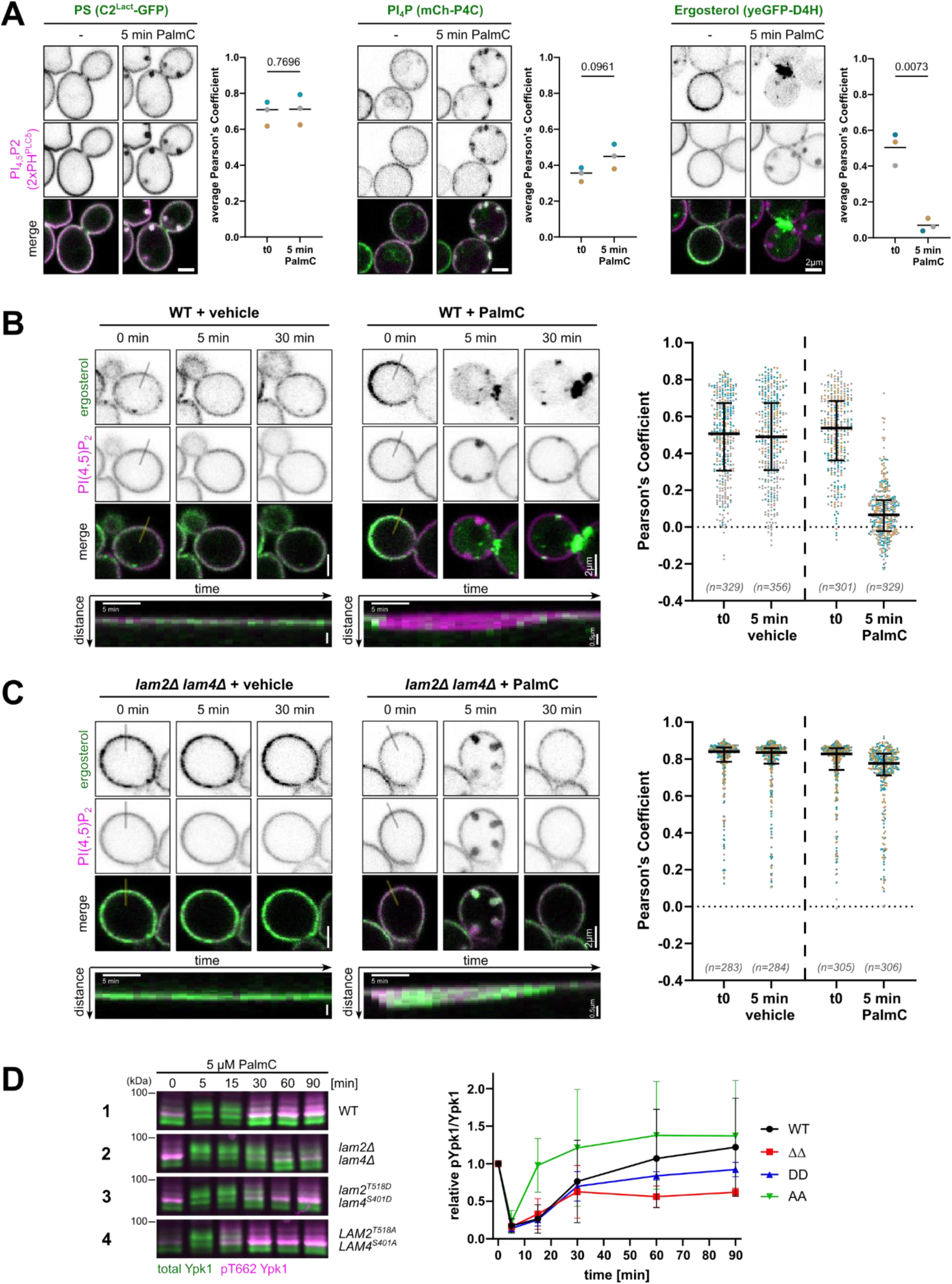
Lam2/4 mediated PM-ergosterol redistribution is required for TORC2 recovery after PalmC. ***A)*** Live-cell fluorescence microscopy of yeast cells expressing a PI(4,5)P2 reporter (GFP- or mCh-2xPH^PLCδ^) along with a phosphatidylserine (PS) reporter (C2^Lact^-GFP), a PI4P reporter (mCh-P4C), or a free ergosterol reporter (yeGFP-D4H). Cells were treated with 5 µM PalmC for 5 minutes, and the colocalization of each reporter with 2xPH^PLCδ^ was quantified (Pearson’s correlation coefficient). Representative images and quantifications (average single-cell values, mean, p-value) are shown. Different colours represent data from independent experiments. ***B, C)*** Live cell fluorescence microscopy of free ergosterol (yeGFP-D4H) and PI(4,5)P2 (mCh-2xPH^PLCδ^) before and after the addition of vehicle or 5 µM PalmC in ***B)*** WT and ***C)*** *lam2Δ lam4Δ* cells. Time-lapse images show the localization of yeGFP-D4H relative to mCh-2xPH^PLCδ^ at the indicated timepoints. Kymographs (bottom panels) depict yeGFP-D4H distribution relative to a mCh-2xPH^PLCδ^-marked PM invagination along the specified line over 30 minutes, at 1-minute intervals. Scatter plots show the colocalization between the two probes before and after 5 minutes of treatment, data points represent individual cells, plotted with median and interquartile range. Different colours represent data from independent experiments. ***D)*** Western blot analysis of TORC2 activity in WT or the indicated mutants after treatment with 5 µM PalmC, as assessed by relative phosphorylation of Ypk1. Mean and SD.

Ergosterol however displays a strikingly different behaviour: PalmC treatment leads to a significant loss of ergosterol signal at the PM, evidenced by a decrease in colocalization with PI(4,5)P_2_ Fig. 2A, S2A (right panel). yeGFP-D4H binds free sterols, i.e. the fraction of unesterified sterols inside membranes that is not in complexes with phospholipids or proteins (Johnson *et al*., 2012; Maekawa and Fairn, 2015). The yeGFP-D4H signal in exponentially growing budding yeast is heterogenous but often enriched in buds and at bud necks (Encinar Del Dedo *et al*., 2021) (see Fig. S2C). Addition of vehicle has no impact on this pattern (Fig. 2B, left panel, Supp. movie 1). Upon PalmC addition, yeGFP-D4H can initially be seen in PM invaginations, similarly enriched as PI(4,5)P_2_ (Fig. S2D), however it rapidly dissociates from the PM, instead showing diffuse cytoplasmic signal and bright intracellular foci (Fig. 2B, right panel, kymograph; S2D, Supp. movie 2 and 2b). In line with our previous observations, two PalmC derivatives (C16-Choline and PHS) which caused PM invaginations and TORC2 inhibition also induced relocation of free PM ergosterol, while an ineffective derivative (C16-Glycerol) did not (Fig. S2E). We conclude that PM ergosterol relocation is a consequence of the PM perturbing properties of small amphipaths.

### PalmC-induced sterol redistribution is Lam2/4 dependent

The relocalisation of PM sterol (and GFP-D4H) to similar intracellular foci has been previously described in fission yeast treated with the Arp inhibitor CK-666, where it was dependent on a StART (steroidogenic acute regulatory protein–related lipid transfer) family sterol transporter (Marek, Vincenzetti and Martin, 2020). In budding yeast, TORC2 has been reported to both regulate (Murley *et al*., 2017) and be regulated by the PM-ER-contact site resident proteins Lam2 and Lam4 (Roelants *et al*., 2018; Topolska *et al*., 2020), which mediate retrograde sterol transport via two StART domains (Gatta *et al*., 2015; Jentsch *et al*., 2018; Tong, Manik and Im, 2018). To test if sterol relocalization upon PalmC treatment is dependent on Lam2/4, we observed sterol behaviour in *lam2Δ lam4Δ* cells. Virtually all mutant cells showed a strong yeGFP-D4H signal at the PM, in agreement with previous work that showed an excess of free PM ergosterol ( Fig. S2F) (Gatta *et al*., 2015). Application of PalmC to *lam2Δ lam4Δ* cells triggered formation of large PM invaginations. In contrast to WT however, we observe that sterols are not removed from the PM and instead continue to colocalize with PI(4,5)P_2_, both inside and outside of PM invaginations (Fig. 2C, Supp. movies 3, 4). This confirms that PalmC-induced sterol depletion from the PM is dependent on Lam2/4.

### Lam2/4 activity is necessary for TORC2 recovery after PalmC

To understand how these changes in sterol mobility affect TORC2 regulation, we followed Ypk1 phosphorylation. In WT cells, TORC2 is rapidly inhibited by PalmC and usually recovers within 60 min (Fig. 2D, line 1). In *lam2Δ lam4Δ* cells, TORC2 inhibition still takes place, but recovery is strongly delayed (Fig. 2D, line 2). This suggests that Lam2/4 are necessary for recovery of TORC2 activity after PalmC. With our phosphospecific antibody, we did not observe an increase in baseline TORC2 activity in *lam2Δ lam4Δ* cells, which had been previously reported by electrophoretic mobility shift (Murley *et al*., 2017).

Ypk1, activated directly by TORC2, inhibits Lam2 and Lam4 through phosphorylation on Thr518 and Ser401, respectively (Roelants *et al*., 2018; Topolska *et al*., 2020). To test if Lam2/4 activity regulation by TORC2-Ypk1 has an impact on TORC2 response after PalmC, we generated CRISPR mutants, substituting these residues either with aspartic acid (DD) or alanine (AA). In cells expressing only phospho-mimetic (DD), and therefore presumably hypoactive Lam2/4, TORC2 activity was delayed with respect to WT, though not as severely as in *lam2Δ lam4Δ* cells (Figure 2D, line 3). In contrast, in cells expressing only phospho-mutant (AA), and therefore hyperactive Lam2/4 (Roelants *et al*., 2018), TORC2 recovery was faster than in WT cells (Fig. 2D, line 4). This suggests that Lam2/4 activity, regulated by TORC2, is important for TORC2 activity recovery after PalmC.

### TORC2 inhibition triggers retrograde transport of PM ergosterol

To pinpoint the specific role of TORC2 activity regulation for sterol trafficking during PalmC treatment, we inhibited TORC2 pharmacologically. In budding yeast, it is possible to render TORC2 sensitive to Rapamycin by deleting the C-terminal part of the TORC2 subunit Avo3. This is paired with a *TOR1-1* allele that forms rapamycin-insensitive TORC1 (Gaubitz *et al*., 2015). Addition of rapamycin, but not DMSO, to *AVO3-ΔCT TOR1-1* cells induced the relocation of ergosterol to internal foci, but only after around 30 minutes (Fig. 3A, Supp. movies 5,6). Even considering that TORC2 inhibition by rapamycin is slower than during PalmC treatment (Gaubitz *et al*., 2015), these sterol kinetics are different from the ones observed after PalmC treatment by an order of magnitude. Thus, while TORC2 activity is clearly necessary to maintain PM ergosterol levels, TORC2 inhibition, and thereby Lam2/4 activation, cannot be the only trigger for PalmC induced sterol removal.

**Fig. 3:**
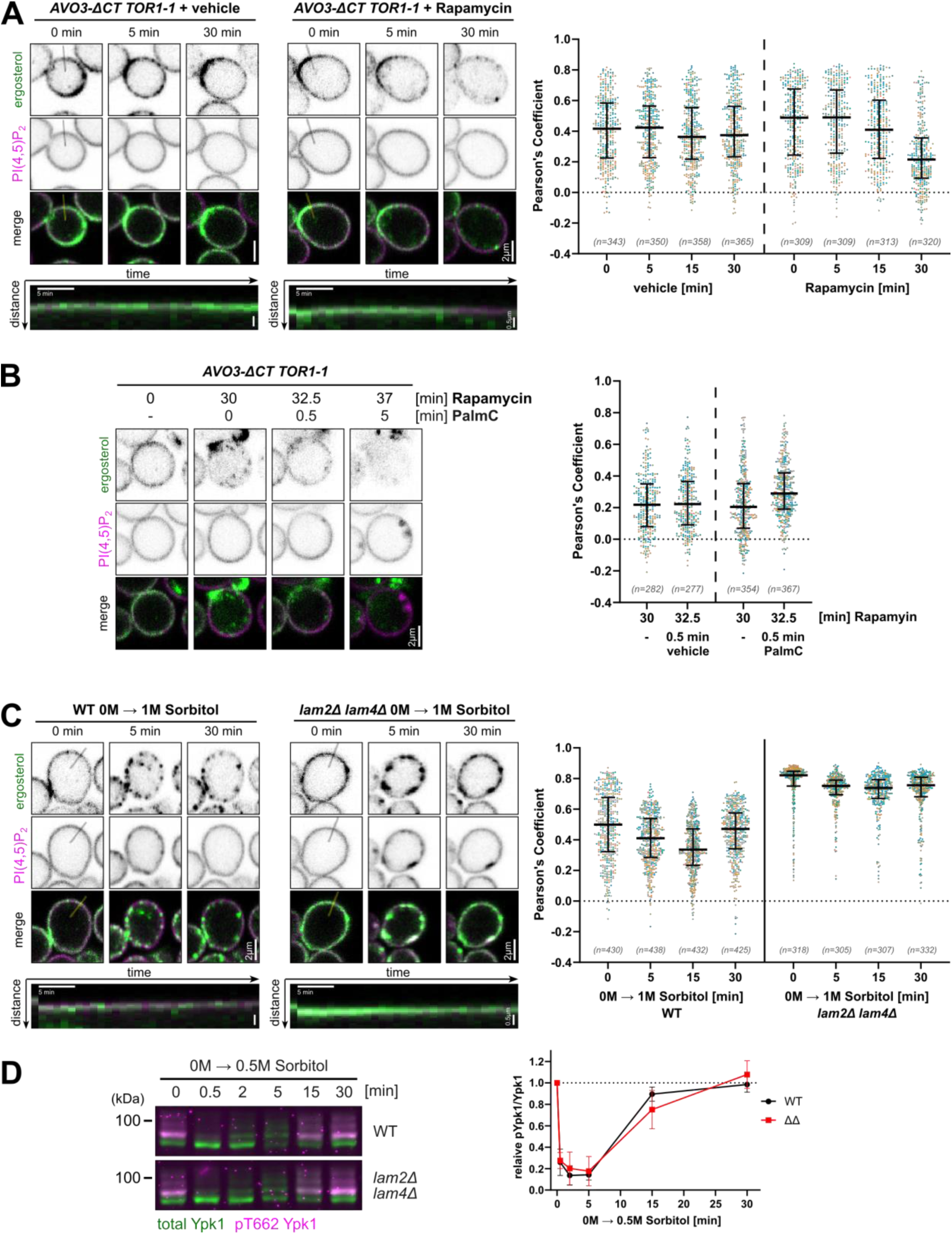
Rapid Ergosterol redistribution is a specific reaction to PalmC. ***A-C)*** Live cell fluorescence microscopy of free ergosterol (yeGFP-D4H) and PI(4,5)P2 (mCh-2xPH^PLCδ^) before and after the addition of ***A)*** vehicle or 200 nM Rapamycin to *AVO3-ΔCT TOR1-1* cells, ***B)*** 200 nm Rapamycin for 30 min, followed by 5 µM PalmC to *AVO3-ΔCT TOR1-1* cells ***C)*** 1 M sorbitol media to WT or *lam2Δ lam4Δ* cells. Time-lapse images show the localization of yeGFP-D4H relative to mCh-2xPH^PLCδ^ at the indicated timepoints. Kymographs (bottom panels, if present) depict yeGFP-D4H distribution relative to a mCh-2xPH^PLCδ^-marked PM invagination along the specified line over 30 minutes, at 1-minute intervals. Scatter plots show the colocalization between the two probes before and at the indicated timepoints post-treatment, data points represent individual cells, plotted with median and interquartile range. Different colours represent data from independent experiments. ***D)*** Western blot analysis of TORC2 activity in WT or *lam2Δ lam4Δ* cells after treatment with 0.5 M sorbitol, as assessed by relative phosphorylation of Ypk1. Mean and SD.

### PalmC transiently increases available ergosterol at the PM

It has been previously shown that small amphipathic molecules render cholesterol molecules in membranes more accessible (Lange *et al*., 2009), presumably by displacing them from microdomains. Thus, we speculated that PalmC integration into the PM might directly stimulate retrograde ergosterol transfer by Lam2/4 by increasing the PM pool of free ergosterol. Indeed, we could observe that immediately after addition of PalmC, the PM signal of yeGFP-D4H increased in some cells (Fig. S3A). However, this effect was clearly visible only in cells that had little to no yeGFP-D4H at the PM before treatment (and thus a PI(4,5)P_2_ colocalization coefficient around 0). Only a very small fraction of cells falls into this category in steady state conditions (see Fig. 2B), rendering this very transient effect difficult to measure across a population in WT cells. Instead, we opted to use Rapamycin (30min) pretreated *AVO3-ΔCT TOR1-1* cells, where PM yeGFP-D4H localization is drastically reduced across the population without direct perturbation of the PM. Addition of PalmC to these cells caused a small but noticeable transient increase of yeGFP-D4H signal at the PM, showing a release of a pool of previously inaccessible ergosterol, which was afterwards rapidly internalized (Fig. 3B, S3B). These results confirm that perturbation of the PM by PalmC leads to the release of a previously inaccessible pool of PM ergosterol.

### Hyperosmotic shock recovery of TORC2 is independent of PM sterol removal

Both PalmC and hyperosmotic shock cause a decrease in membrane tension and TORC2 inhibition (Riggi *et al*., 2018). Since artificial extraction of sterols from membranes has previously been shown to increase membrane tension (Biswas *et al*., 2019; Cox *et al*., 2021), we wondered if this free sterol-TORC2-Ypk1-Lam2/4 feedback loop is a general response that serves to restore decreases in membrane tension by removing lipids – and thus membrane area – from the PM. Unlike PalmC, hyperosmotic shock doesn’t directly alter PM composition by addition of exogenous lipid. If sterol influx from the PM is triggered by a decrease in membrane tension, and serves to alleviate this stress, we would expect that TORC2 recovery is also dependent on sterol internalization during hyperosmotic shock.

Both in WT and in *lam2Δ lam4Δ* cells, yeGFP-D4H formed clusters at the PM upon hyperosmotic shock with 1M sorbitol, which often overlapped with the shallow PM invaginations observed with mCh-PH^PLCδ^. After around 10-15 min we observed a slow internalisation of a minor fraction of yeGFP-D4H signal to internal foci in WT, but not in *lam2Δ lam4Δ* cells (Fig. 3C, Supp. movies 7,8). However, the vast majority of yeGFP-D4H signal remained at the PM. Furthermore, Lam2/4 were not instrumental in TORC2 recovery from hyperosmotic shock, which happened at the same rate in *lam2Δ lam4Δ* and WT cells (Fig. 3D, S3C). This suggests that PM ergosterol removal is a specific response to mitigate the PM perturbing effects of small amphipaths like PalmC, rather than a general response to a decrease in PM tension.

## Discussion

We identified a family of small amphipathic molecules that intercalate into the plasma membrane, causing large PM invaginations and TORC2 inhibition, similar to PalmC (Riggi *et al*., 2018). Further characterization revealed that small amphipath-induced PM perturbation leads to rapid ergosterol relocation dependent on PM-ER sterol transporters Lam2/4, which are important for TORC2 activity recovery. The mechanistic framework for this feedback loop is summarized in Fig. 4.

**Fig. 4:**
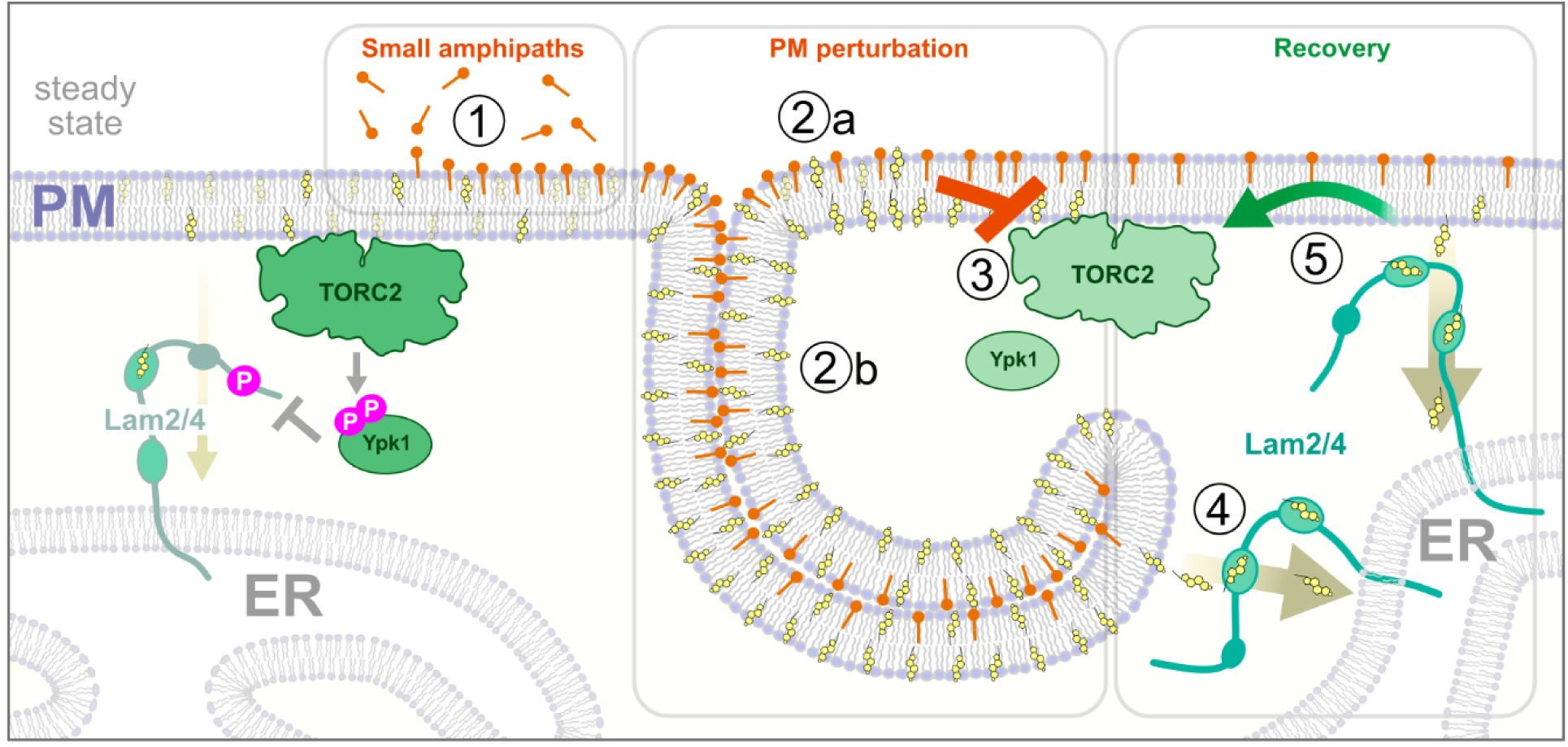
Model for the mechanism of small amphipaths. During steady state, the majority of PM ergosterol is biochemically inaccessible, and homeostatic signalling by TORC2-Ypk1 negatively regulates Lam2/4 activity. **①** Large amounts of small amphipaths intercalate into the PM. **②a** Small amphipaths release sterols from lipid complexes **②b** Formation of giant PM invaginations. **③** TORC2 activity is inhibited by the PM perturbation. **④** Lam2/4 mediate retrograde PM-ER ergosterol transport. **⑤** A decrease in free PM ergosterol recovers TORC2 activity.

PalmC has been previously shown to cause a perturbation of PM biophysical properties, i.e. PM tension and lipid order. The observation that PalmC also affects ATP-depleted cells and giant unilamellar vesicles (GUVs) suggested that it acts passively by direct intercalation into the PM (Riggi *et al*., 2018). Our results corroborate that notion: the fact that the effect of small amphipaths is determined by the total dose per cell suggests that large amounts need to intercalate into the PM to physically perturb it (Fig. 4, ①), and our SAR screen showed that the effect of PalmC is a consequence of its physicochemical properties rather than specific chemistry. Derivatives with longer carbon chains, and thus increased hydrophobicity and membrane partitioning (Requero, Goni and Alonso, 1995; Ho, Duclos and Hamilton, 2002) caused stronger effects. This confirms that their primary action is to insert into the PM, as opposed to directly binding to TORC2. Furthermore, comparison of the naturally occurring sphingolipid precursor D-*erythro-*DHS and the non-metabolizable L-*erythro-*DHS (Watanabe *et al*., 2002) showed that their effect as TORC2-inhibitors is virtually indistinguishable. This observation strongly speaks against a metabolic component of the rapid effects of PalmC. Our findings could extend to physiological or pharmacological molecules with similar properties to PalmC. For instance, Alkylphospholipid analogs (Fei *et al*., 2023). have been shown to down-regulate TORC2 signalling (Gills and Dennis, 2009; Nomura and Inoue, 2024) and relocalize sterols (Zaremberg *et al*., 2005).

Intercalation of small amphipaths triggers the appearance of large PM invaginations (Fig 4, ②b), correlating with TORC2 inhibition. Constitutive PM invaginations in yeast have previously been associated with PI(4,5)P_2_ or TORC2 misregulation, but their exact role in TORC2 inhibition remains unclear (Singer-Krüger *et al*., 1998; Stefan, Audhya and Emr, 2002; Walther *et al*., 2006; Berchtold *et al*., 2012; Rodríguez-Escudero *et al*., 2018; Sakata *et al*., 2022). We previously proposed that PI(4,5)P2 enrichment in PM invaginations was important for PalmC-induced TORC2 inactivation, using the heat sensitive PI(4,5)P2 kinase allele *mss4^ts^ -* a rather blunt tool (Riggi *et al*., 2018). Wishing to challenge our previous hypothesis, we expanded our investigation to monitor the behaviors of other membrane lipids, leading to a new model: small amphipaths act as surfactants that perturb membrane lipid organization, leading first to an increase of free ergosterol at the PM (Fig. 4, ②a). This is in agreement with previous studies reporting that small amphipaths dissolve membrane domains and displace sterols from lipid complexes (Zaremberg *et al*., 2005; Lange *et al*., 2009). PM sterols are now more available for D4H- and Lam2/4-binding. This likely already amplifies Lam2/4 activity, as sterol redistribution over non-vesicular sterol transporters is thought to be majorly regulated by the availability of sterol itself (Lange, Ye and Steck, 2004; Lange and Steck, 2020; Koh *et al*., 2023; Steck and Lange, 2023). While membrane phase behaviour, and thereby free sterols, can also be influenced by hyperosmotic shock (García-Sáez, Chiantia and Schwille, 2007; Hamada *et al*., 2011; Roffay *et al*., 2021; Shimokawa and Hamada, 2023; Torra, Campelo and Garcia-Parajo, 2024) and TORC2 inhibition (Roelants *et al*., 2017; Riggi *et al*., 2019, 2019; Thorner, 2022), our results suggest that small amphipaths have a much stronger effect. In fact, even after TORC2-induced PM sterol removal, PalmC still triggers the release of a previously inaccessible pool of ergosterol at the PM. The rapid increase of free sterol at the PM correlates temporally with TORC2 inhibition (Fig. 4, ③), and PM sterol removal by Lam2/4 (Fig. 4, ④), further stimulated by the inhibition of TORC2 (Roelants *et al*., 2018; Topolska *et al*., 2020), is necessary to recover TORC2 activity after small amphipath treatment (Fig 5, ⑤) – but not after hyperosmotic shock. It is unclear if the excess of free sterols itself is part of the inhibitory signal to TORC2 or merely a symptom of the TORC2-inhibiting PM perturbation. However, previous studies have suggested that TORC2 is sensitive to changes in PM domains (Berchtold *et al*., 2012; Riggi *et al*., 2018; Sakata *et al*., 2022), and sterols are major modulators of membrane domain formation (Wolf *et al*., 2001; Bacia, Schwille and Kurzchalia, 2005; Dufourc, 2008; Levental and Wang, 2020). For example, in the eisosomal microdomain scaffolded by Pil1/Lsp1, which sequesters the TORC2 activators Slm1/2 (Berchtold *et al*., 2012), ergosterol is mobilized upon stretch of the scaffold. (Kefauver *et al*., 2024), suggesting it may also have a role in the TORC2 activation mechanism.

Collectively, our data describe a feedback loop wherein an increase in free sterols leads to TORC2 inhibition, which subsequently stimulates free sterol removal to alleviate this PM stress. We propose that TORC2 monitors the integrity of PM domains or phases as a global factor, possibly by sensing changes in free sterol levels. It would be interesting to see if retrograde sterol transport plays a role in TORC2 recovery during other, more physiological stresses, such as e.g. heatshock, which like PalmC (Kobayashi *et al*., 1989; Riggi *et al*., 2018) is expected to affect membrane fluidity and phase behaviour (Beney and Gervais, 2001; Los and Murata, 2004; Blicher *et al*., 2009; Fonseca *et al*., 2019).

### Limitations of the study

The behaviour of membrane lipids *in vivo* is notoriously hard to observe. In this study, we made use of fluorescent lipid associated reporters (FLAREs) as powerful tools to study dynamic lipid behaviour *in vivo.* The observations made with these FLAREs are however subject to the limitations of these biosensors, i.e. they compete with endogenous lipid interactors which 1) potentially changes lipid and binding partner behaviour, 2) allows them to only interact with certain (free) pools of the detected lipids. This is especially relevant for the D4H probe, which only binds to membranes containing sterols above a certain threshold (Johnson *et al*., 2012; Maekawa, Yang and Fairn, 2016). More research is needed to distinguish between chronic effects of Lam deletion/activity changes (and the resulting difference in PM sterol levels) and the effects of acute Lam2/4 stimulation, and to understand the mechanistic link – if existent – between free PM sterols and TORC2 inhibition.

## Material and Methods

### Yeast cell culture and treatments

Used yeast strains and plasmids are listed in Supp. Tables 1 & 2. Yeast strains were generally generated using classical recombination techniques. Strains with point mutations were generated using CRISPR-Cas9-based methods.

Yeast cells were routinely grown at 30°C in synthetic media (SC) buffered at pH 6.2 with 0.1 M Sorensen Buffer and supplemented with 2% glucose and appropriate amino acids and nucleobases to maintain plasmids. For microscopy experiments, cells were grown in low fluorescence media. All experiments were performed in logarithmically growing cells.

All treatment substances and substance stocks are listed in Supp. table 3. For small amphipath treatments, cells were usually grown to OD_600nm_ 0.6-0.7 and treated with the indicated substances at 5 µM, or an equivalent volume of vehicle (DMSO or Methanol). To test the impact of culture density on PalmC effect, cells were pelleted by centrifugation, resuspended to OD_600nm_ 0.1 or 2 in fresh prewarmed media, and incubated for 15-20 min at 30°C to recover before administering 5 µM PalmC. To negate the effect size variation caused by offsets in the small amphipath/cell ratio, all experiments with small amphipaths were performed at the same OD_600nm_ for compared conditions. For hyperosmotic shocks, cultures were diluted with prewarmed media containing 2M sorbitol to a final concentration of 0.5M or 1M sorbitol. Samples were harvested at the indicated timepoints and processed as described below.

### Mammalian cell culture and treatments

HBEC3-KT (human bronchial epithelium immortalized with hTERT) cells were kindly provided by Prof. Georgia Konstantinidou. They were cultured in Keratinocyte serum-free medium (SFM) supplemented with human recombinant epidermal growth factor (rEGF) and bovine pituitary extract (17005042, Life Technologies) at 37°C with 5% CO2. Cells were authenticated by Microsynth and tested negative for mycoplasma by GATC Biotech. For WB experiments, mammalian cells were seeded on 6-well plates a day prior to the experiment. Treatments were performed by exchanging culture media for prewarmed media supplemented with the indicated substances. At the indicated timepoints, cells were rinsed with ice-cold PBS quickly before lysis (described below).

### WB sample preparation and detection of phosphoproteins on WB

Yeast culture aliquots were processed according to standard TCA-urea extraction procedures. In brief, culture aliquots were incubated with 5% TCA on ice for at least 10 min, before pelleting and drying cells with ice cold acetone. Cells were lysed by bead beating in a urea buffer (25 mM TRIS pH6.8, 6M Urea, 1% SDS), and boiled for 5 min. Denatured lysates were mixed with 2x sample buffer (125 mM TRIS pH6.8, 20 vol% Glycerol, 2% SDS, 0.02% Bromphenol Blue, 200mM DTT), and boiled again before analysing them via WB. Mammalian cells were lysed with a lysis buffer (40mM HEPES, 10mM Na-PPi, 10mM Na-β-glycerophosphate, 4mM EDTA 1% Triton X-100, pH 7.4) supplemented with 1× Halt Protease & Phosphatase Inhibitor Cocktail (Thermo Fisher Scientific) and kept at -20°C until further analysis. A Pierce BCA Protein Assay (Thermo Fisher Scientific) was used to determine protein concentration in the samples.15-50 µg of total protein of each sample was used for further analysis. The 5x sample buffer (312.5mM Tris, 10% SDS, 50% glycerol, 25% β-mercaptoethanol, 0.1% bromophenol blue, pH 6.8) was added and samples were denatured at 95°C for 5min.

Protein lysates were separated on 7.5% (Ypk1, Sch9) or 10% (Ypk1, Akt) SDS-page gels and blotted onto nitrocellulose membranes using the iBlot2 Gel Transfer System (Thermo Fisher Scientific). Membranes were blocked with BSA, incubated with primary antibodies overnight at 4 °C, washed and incubated with secondary antibodies for 1 hour at room temperature, using PBS-(yeast WBs) or TBS (yeast and mammalian WBs) -based buffers. Used antibodies are listed in Supp. Table 4. Membranes were developed using an Odyssey imaging system (LI-COR) and signal intensities were quantified using FIJI (ImageJ 1.54f), calculations were performed in Microsoft Excel, and data was plotted using GraphPad Prism (10.2.3).

### Yeast membrane extraction

1L culture was harvested by centrifugation (5min, 5000g), and resuspended in 80mL 0.4M sucrose Buffer A - 25mM imidazole (pH 7.0) containing 2.5µg/mL pepstatin A (Sigma, St. Louis, MO), and a 1/100 dilution of complete protease inhibitor cocktail (Roche Molecular Biochemicals, Basel, Switzerland) from a stock in H2O (two tablets dissolved in 2mL H2O). After centrifugation (5000g for 10min), the pellet was covered by twice its volume of glass beads followed by just enough 0.4 M sucrose in buffer A to cover the cells and glass beads. The solution was vortexed 3 times for 2min, then diluted 3 times in 0.4 M sucrose in buffer A and centrifuged at 530g for 20min. The supernatant was recentrifuged at 20000g for 30min. The pellet from this last step was then resuspended in 2mL buffer A, and 1mL aliquots were loaded onto discontinuous sucrose gradients and centrifuged overnight (14h) at 80000g and 4°C. Membranes banding at the 2.25/1.65M sucrose interfaced were collected, diluted 4 times in buffer A and pelleted by centrifugation at 30000g and 4°C for 40min. They were stored and -20°C.

### Metabolite extraction

Frozen cell lysates and membrane fraction from yeast were pre-extracted and homogenized by the addition of 1mL of MeOH:H_2_O (4:1), in the Cryolys Precellys 24 sample Homogenizer (2 x 20 seconds at 10000 rpm, Bertin Technologies, Rockville, MD, US) with ceramic beads. The bead beater was air-cooled down at a flow rate of 110 L/min at 6 bars. Homogenized extracts were centrifuged for 15min at 4000g at 4°C (Hermle, Gosheim, Germany). The supernatant (metabolite extract) was collected and evaporated to dryness in a vacuum concentrator (LabConco, Missouri, US). Dry cell and plasma membrane extracts were reconstituted in 300μL of MeOH:H_2_O (4:1). For absolute quantification of PalmC the samples were prepared by mixing an aliquot (5μL) of reconstituted extract with 250μL of the ice-cold internal standard solution (in 100% methanol) and 45μL of 0.1% formic acid in water. These mixtures were centrifuged and supernatant was injected for LC-HRMS analysis in positive ionization mode.

### Protein quantification for normalization of PalmC levels

Following the metabolite extraction, the excess of organic solvent remaining on the top of precipitated protein pellets was evaporated and the protein pellets were resuspended in the lysis buffer containing: 20mM Tris-HCl (pH 7.5), 4M guanidine hydrochloride, 150mM NaCl, 1mM Na_2_EDTA, 1mM EGTA, 1% Triton, 2.5mM sodium pyrophosphate, 1mM beta-glycerophosphate, 1mMNa3VO4, 1μg/ml leupeptin; using brief probe-sonication (5 pulses x 5 sec). BCA Protein Assay Kit (Thermo Scientific, Massachusetts, US) was used to measure (A562nm) total protein concentration (Hidex, Turku, Finland).

### Hydrophilic Interaction Liquid Chromatography coupled to high resolution mass spectrometry (HILIC-HRMS)

For PalmC quantification, the prepared extracts were analyzed by Hydrophilic Interaction Liquid Chromatography coupled to high resolution mass spectrometry (HILIC - HRMS) in positive ionization mode. A Vanquish Horizon (Thermo Fisher Scientific) ultra-high performance liquid chromatography (UHPLC) system coupled to Q Exactive™ Focus interfaced with a HESI source was used for the quantification of amino acids. Chromatographic separation was carried out using an Acquity BEH Amide (1.7 μm, 100 mm × 2.1 mm I.D.) column (Waters, Massachusetts, US). The mobile phase was composed of A = 20mM ammonium formate and 0.1 % formic acid in water and B = 0.1 % formic acid in ACN.

The gradient elution started at 95% B (0-2 min) decreasing to 65% B (2 min – 14 min), reaching 50% B at 16 min, followed by an isocratic step (16min – 18min) and 4min post-run for column re-equilibration. The flow rate was 400μL/min, column temperature 25°C and the sample injection volume was 2μl. HESI source conditions operating in positive mode were set as follows; sheath gas flow at 60, aux gas flow rate at 20, sweep gas flow rate at 2, spray voltage at +3kV, capillary temperature at 300°C, s-lens RF level at 60 and aux gas heater temperature at 300°C. Full scan HRMS acquisition mode (m/z 50−750) was used with the following MS acquisition parameters; resolution at 70,000 FWHM, 1 microscan, 1e6 AGC and 100ms as maximum inject time.

Data was processed using Xcalibur (version 4.1, Thermo Fischer Scientific). For absolute quantification, calibration curve and the stable isotope-labeled internal standard (ISTD – PalmC-d9) was used to determine the response factor. Linearity of the standard curves was evaluated using a 9-point range; in addition, peak area integration was manually curated and corrected where necessary. Concentration of PalmC was corrected for the ratio of peak intensity (peak area) between the analyte and the ISTD, to account for matrix effects. Measured metabolite quantities were normalized to protein content and expressed in nmoles/mg of protein, to normalize for sample amount differences.

### SAR screen live-cell fluorescent microscopy

For live cell imaging, TB50A WT cells expressing GFP-2xPH^PLCδ^ from plasmid were mounted onto Concanavalin A-coated VI 0.4 µ-Slides (Ibidi) primed with media (except LCBs). For substance treatment, media were removed and exchanged with small amphipath containing media at RT. Images of the equatorial plane were taken at the indicated timepoints on a Leica TCS SP5 gatedSTED CW microscope with a 63.0×1.40 oil immersion objective and LAS AF (Ver. 2.7.3.9723) software. The effects of LCBs were extremely variable under these conditions, thus D-DHS and PHS were instead added to 5 µM to shaking yeast cultures, cells were concentrated by brief centrifugation at the indicated timepoints, mounted onto glass slides, and imaged using a Zeiss LSM 800 microscope with a 63×1.40 oil immersion objective and Zen Blue (Ver. 2.6) software. Images were processed in FIJI (ImageJ 1.54f).

### SAR screen PM invagination quantification by automated confocal microscopy

The screen was performed in collaboration with the ACCESS facility Geneva. TB50A WT cells expressing mCherry-2xPH^PLCδ^ from the HIS3 locus were grown logarithmically and set to OD_600nm_ 0.6. The culture was split into subcultures of same volume, agitating at 30°C, and substances were added to 5 µM from prewarmed stocks. At indicated timepoints, culture aliquots were taken and mixed 1:2 with Paraformaldehyde 3% Glutaraldehyde 0.35% In 0.1M Sodium Cacodylate Buffer pH 7.4 (Electron Microscopy Sciences). Cells were fixed for 30-60 min at RT and diluted in low fluorescence media. C14-C invaginations could not be preserved in fixed cells. Images of the equatorial plane were taken on a Molecular Devices™ ImageXpress Micro HTai C® High-Content Imaging System equipped with a 60x 1.25 NA water immersion objective. Single cell and PM invagination segmentation and analysis was performed with custom module editor from MetaXpress software (Ver 6.7.0.211). Briefly, individual yeasts were segmented on a log filtered transmitted light image. The PM invaginations were segmented on the mCherry-2xPH image after top hat deconvolution. The area sum of PM invaginations was normalized to the cell area and averaged for each condition and timepoint. These average values where then normalized to the value determined in vehicle treated cells at the same timepoint for each experiment. Calculations were performed in Microsoft Excel, and data was plotted using GraphPad Prism (10.2.3).

### Live cell microscopy of Fluorescent Lipid Associated Reporters

WT or mutant yeast cells expressing FLAREs from genomic loci (mCherry-2xPH^PLCδ^ + GFP-D4H) or from plasmids (all other combinations) were grown to mid log phase in low fluorescence media and imaged at room temperature on a Leica STELLARIS 8 FALCON FLIM Microscope with a 63×1.4 Oil immersion objective and LAS X (Ver. 4.6.1.27508) software. For vehicle, PalmC, Rapamycin and 1 M sorbitol treatment, cells were mounted onto Concanavalin A-coated VI 0.4 µ-Slides (Ibidi) primed with media. The channel was gently flushed with media once to obtain a single layer of cells. Several positions were selected proximal to the “+” channel opening, and equatorial plane images of the same positions/cells were taken at t0, and at 1 min intervals or at the indicated timepoints post treatment. For treatments, media supplemented with indicated substances were added to the “+” channel opening while the excess was absorbed with a paper tissue inserted into the “-“ channel end. To test the effects of C16-Glycerol, C16-Choline and PHS on yeGFP-D4H localization, 5 µM of these substances were instead added to mid log phase liquid cultures to avoid the previously observed substance absorption to the slide. Before the indicated timepoint, cells were concentrated by short centrifugation and mounted onto high precision glass coverslips for imaging. Images were processed using FIJI (ImageJ 1.54f). Before the generation of kymographs, timelapses of selected cells were stabilized using the Image Stabilizer plugin for ImageJ (Li, 2008).

### FLARE colocalization analysis

In flow chambers, we typically observed different and more variable small amphipath kinetics than in liquid culture. We decided to use the 5 min timepoint for quantifications with PalmC, as it generally displayed a robust phenotype. To calculate colocalization between 2xPH^PLCδ^ and other FLAREs, cells were first segmented using Cellpose (Stringer *et al*., 2021). Cells that were intersected by image borders were excluded. Pearson’s correlation coefficient (PCC) as a score for colocalization between different FLAREs and 2xPH^PLCδ^ in single cells was calculated using the BIOP version of the JACoP plugin (Bolte and Cordelières, 2006). Data were analysed and plotted using GraphPad Prism (10.2.3).

### FLARE sorting coefficient analysis

PM invagination sorting coefficients were calculated for each flare using 2xPH^PLCδ^ as reference with the following equation:

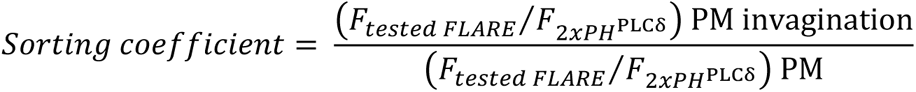

where *F*_*tested FLARE*_ and 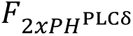 are the background-subtracted fluorescence densities integrated from the plot profiles of PM invaginations with unsaturated signal or an adjacent non-invaginated PM patch in the same cell. Only one invagination was measured in each cell.

Microscopy images were visualized, and data extracted using FIJI (ImageJ 1.54f). Data was analysed using Origin Pro 2024 (OriginLab Corp.) and plots were generated using GraphPad Prism (10.2.3).

### Statistics and reproducibility

Western blot results for TORC2 and TORC1 activity (pT662 Ypk1 and pS758 Sch9, respectively) are presented as mean values and SD from at least three independent replicates. SAR live cell microscopy experiments were performed three times for each compound with similar results, and representative cells are shown. The SAR high content microscopy screen was performed three times and the average, vehicle-normalized value for each condition is plotted together with the mean and SD of all three experiments. To assess differences in FLARE colocalization upon PalmC treatment in WT cells, the experiment was repeated three times, analysed in single cells, and statistical significance was calculated from the mean Pearson’s correlation coefficients using unpaired t-test. Observations of D4H behaviour under different conditions were repeated at least three times with similar results. Single cell scatter plots represent pooled datasets from 2-3 experiments (colour coded), to analyse around 300 cells per condition (n counts are indicated in the graphs); plotted with median and interquartile range (bottom 25% to top 75% quartile). Sorting coefficient values were determined from one invagination/cell; the values from 2-3 independent experiments (colour coded, n counts indicated in the graphs) are plotted together with median (line), interquartile range (bottom 25% to top 75% quartile, box), and range (whiskers).

## Author Contributions

M.G.T, P.N., A.R. and R.L. designed the project. M.G.T., B.K., P.N., and J.K. designed and/or performed experiments in yeast. P.N. performed experiments in mammalian cells. V.M. acquired, processed and quantified images for the high content microscopy screen. M.G.T., P.N., R.L. and A.R wrote the manuscript. A.R. and R.L. supervised the project. All authors discussed the results and commented on the manuscript.

## Acknowledgements

RL acknowledges support from the Swiss National Science Foundation the Canton of Geneva and the European Research Council (ERC AdG TENDO). A.R. acknowledges funding from the Swiss National Fund for Research grant numbers #CRSII5_189996 and #310030_200793 and the European Research Council Synergy grant number #951324-R2-TENSION. RL and AR further acknowledge support from the SNSF NCCR Chemical Biology.

This work used the Photonics imaging platform, and the ACCESS platform at the University of Geneva. Metabolite extraction, protein quantification and HILIC-HRMS for PalmC quantification were performed and evaluated at the Metabolomics platform at the University of Lausanne. We thank Margot Riggi for collecting the yeast cell lysates and membrane fractions for PalmC quantification, Kerstin Hinterndorfer for generating the pRS406-mCh-2xPH(PLCδ) plasmid and the original mCh-2xPH(PLCδ) yeast strain, Ariane Bergmann for generating CRISPR mutants, and Paraskevi Linardou for technical support in generating yeast strains. We thank Marko Kaksonen, Sophie Martin and Christopher Stefan labs for sharing plasmids and/or materials, and, together with Loewith and Roux labs, for helpful discussions.

