## Supplementary Figures for "TORC2-regulated sterol redistribution mediates recovery from membrane perturbation by small amphipathic molecules"

### Supplementary information

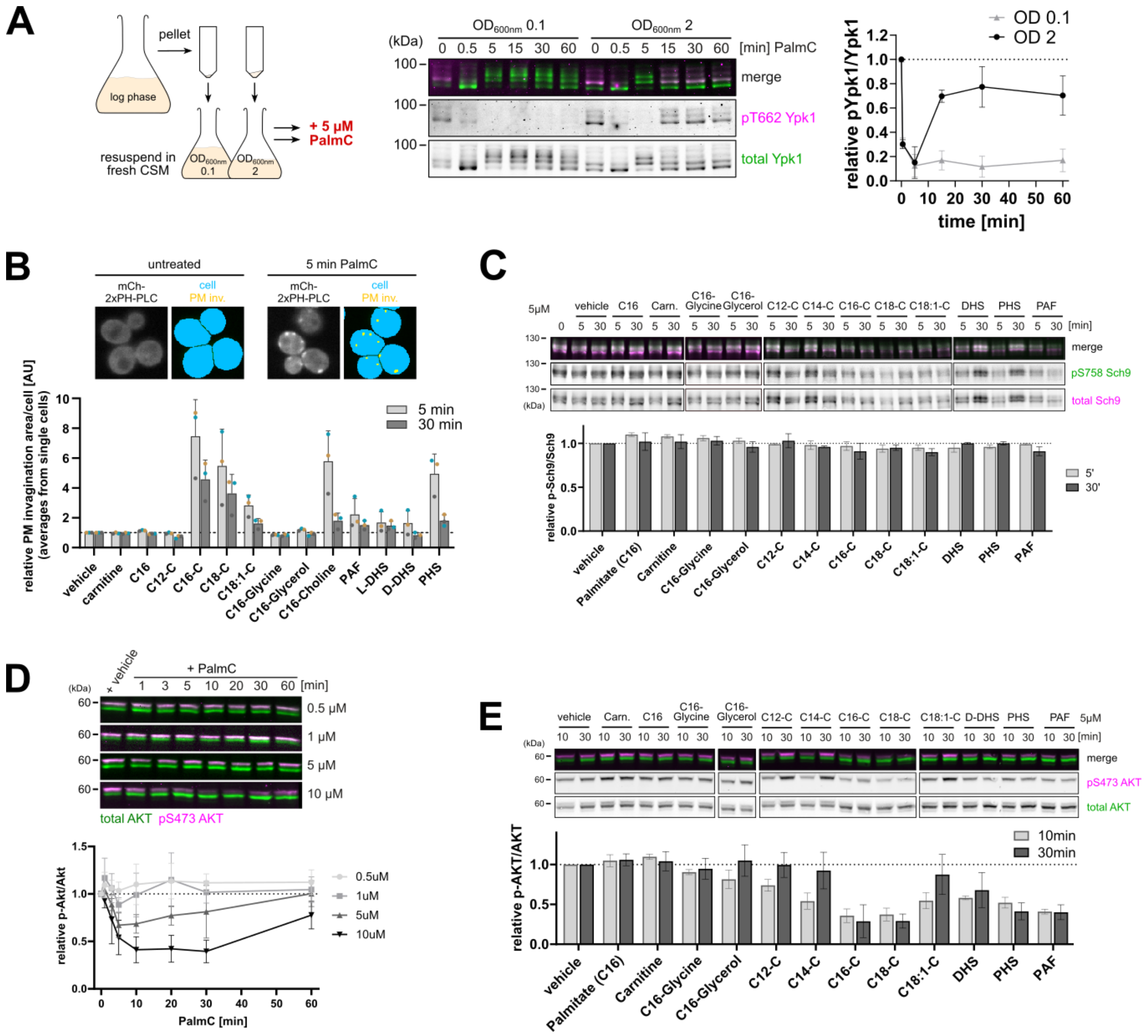


**Supplementary Fig. 1.**

***A)*** Western blot analysis of the effect of different yeast culture densities on PalmC-induced inhibition of TORC2. WT cells were pelleted and resuspended in fresh media to either OD 0.1 or OD 2 before treatment with 5 µM PalmC, and TORC2 activity was assessed by relative phosphorylation of Ypk1. Mean and SD.

***B)*** PM invagination screen in fixed WT cells expressing mCh-2xPH$^{PLC\delta}$. Cells were treated with the indicated substances at 5 µM, and samples were taken at the indicated timepoints and fixed. Representative images and segmentations of untreated cells and cells treated with PalmC for 5 min are shown. The plot shows the relative cell area fraction occupied by PM invaginations under different treatments. Determined values for each condition were normalized to the same vehicle timepoint for each experiment. Mean and SD.

***C)*** Western blot analysis showing the effect of different PalmC derivatives on TORC1 activity. WT cells were treated with indicated substances at 5 µM for 5 or 30 min, and TORC1 activity was assessed by relative phosphorylation of Sch9. Mean and SD.

***D,E)*** Western blot analysis of mTORC2 activity in HBEC3-KT cells. Cells were treated with ***D)*** different concentrations of PalmC, or ***E)*** with the indicated PalmC derivatives at 5 μM, and mTORC2 activity was assessed by relative phosphorylation of AKT. Mean and SD.

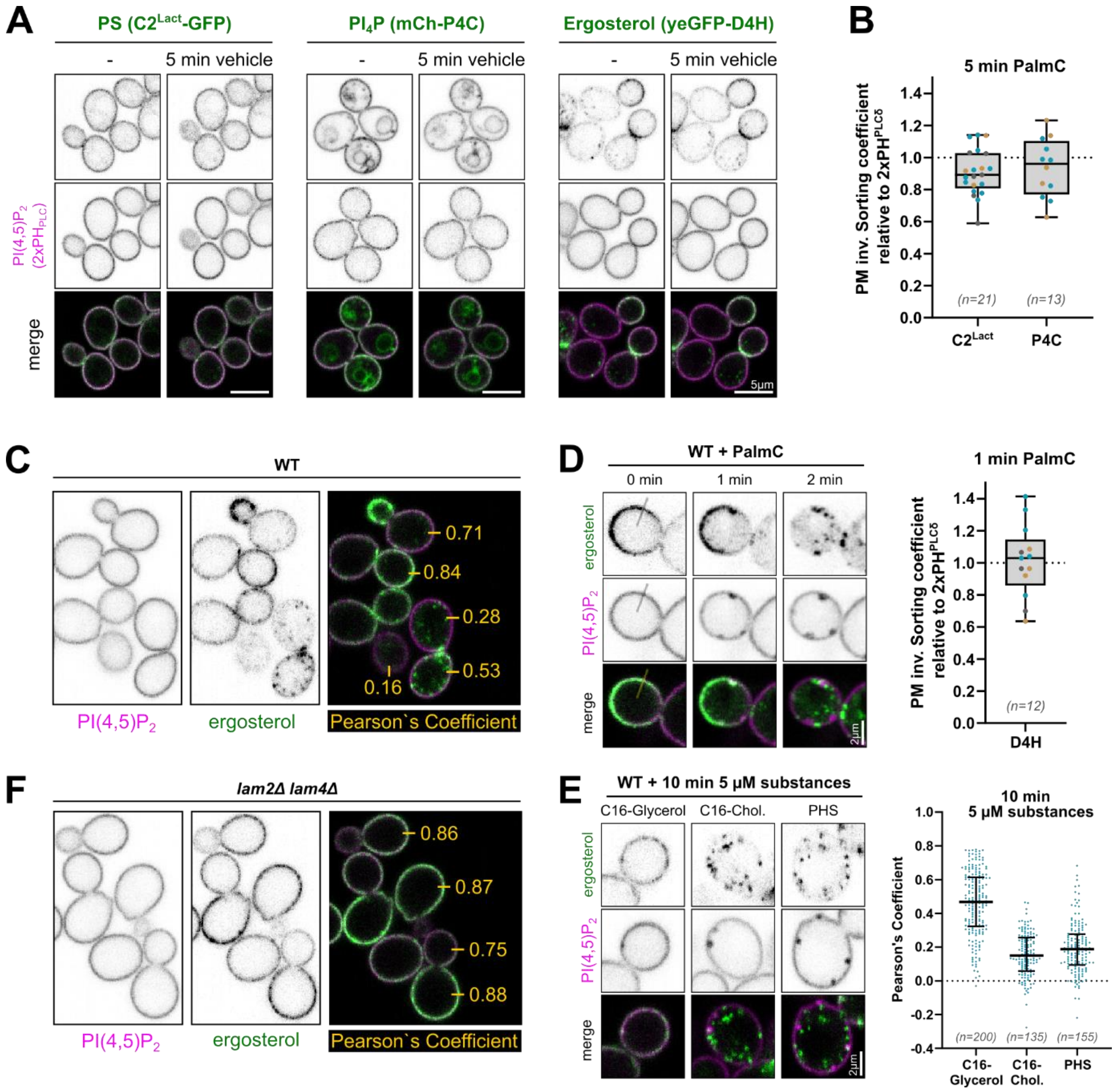


**Supplementary Fig. 2.**

***A)*** Live-cell fluorescence microscopy of yeast cells expressing a PI(4,5)P2 reporter (GFP- or mCh-2xPH$^{PLC\delta}$) along with a phosphatidylserine (PS) reporter (C2$^{Lact}$-GFP), a PI4P reporter (mCh-P4C), or a free ergosterol reporter (yeGFP-D4H). Cells were treated with vehicle for 5 minutes.

***B)*** Relative enrichment (sorting coefficients) of C2$^{Lact}$-GFP and mCh-P4C in PM invaginations after 5 min 5 µM PalmC treatment, calculated using 2xPH$^{PLC\delta}$ as a reference. Single values from independent experiments (colour coded) are plotted together with median, interquartile range, and range.

***C, F)*** Representative image showing variability of yeGFP-D4H distribution and corresponding Pearson`s Correlation Coefficients as measure of colocalisation with mCh-2xPH$^{PLC\delta}$ across the population in ***C)*** WT and ***F)*** *lam2Δ lam4Δ* cells.

***D)*** Live cell fluorescence microscopy of free ergosterol (yeGFP-D4H) and PI(4,5)P2 (mCh-2xPH$^{PLC\delta}$) before, and at indicated timepoints after the addition 5 µM PalmC. The box plot shows relative enrichment (sorting coefficient) of yeGFP-D4H in PM invaginations after 1 min 5 µM PalmC treatment, calculated using 2xPH$^{PLC\delta}$ as a reference. Single values from independent experiments (colour coded) are plotted together with median, interquartile range, and range.

***E)*** Live cell fluorescence microscopy of free ergosterol (yeGFP-D4H) and PI(4,5)P2 (mCh-2xPH$^{PLC\delta}$) in yeast cells treated with indicated PalmC derivatives at 5 µM for 10 minutes. Scatter plots show the colocalization between the two probes, data points represent individual cells, plotted with median and interquartile range. Values from one representative experiment are shown.

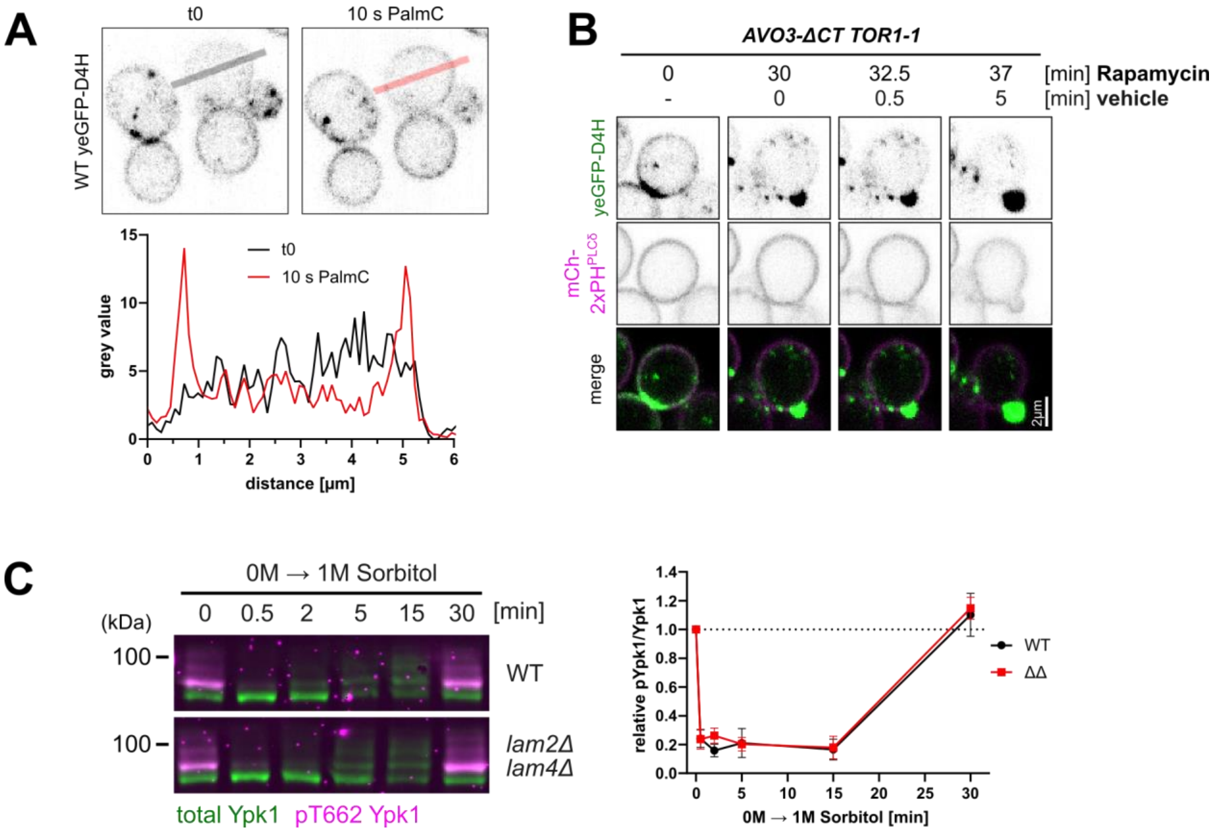


**Supplementary Fig. 3.**

***A)*** Live cell fluorescence microscopy of free ergosterol (yeGFP-D4H) in WT cells before and 10s after 5 µM PalmC addition to WT cells. The line plot represents the average signal intensity of yeGFP-D4H along the specified line.

***B)*** Live cell fluorescence microscopy of free ergosterol (yeGFP-D4H) and PI(4,5)P2 (mCh-2xPH$^{PLC\delta}$) in *AVO3-ΔCT TOR1-1* cells before and after 30 min pretreatment with 200 nm Rapamycin, followed by addition of vehicle.

***C)*** Western blot analysis of TORC2 activity in WT or *lam2Δ lam4Δ* cells after treatment with 1 M sorbitol, as assessed by relative phosphorylation of Ypk1. Mean and SD.

| Name | genotype | source |
|---|---|---|
| TB50a | *MATa leu2-3,112 ura3-52 rme1 trp1 his3* | Loewith lab |
| TB50α | *MATα leu2-3,112 ura3-52 rme1 trp1 his3* | PMID:26028537 |
| BK2-8 | TB50α *lam2::Nat lam4::Nat* | This study |
| PN001 | TB50a *LAM2-T518A LAM4-S401A* | This study |
| PN002 | TB50α *LAM2-T518D LAM4-S401D* | This study |
| MTY002 | TB50a + *pRS426-GFP-2xPH(PLCδ)* | This study |
| BK3 | TB50a + *pRS414-pTdh3-mCherry-2xPH(PLCδ)* + *pRS416-LactC2-GFP* | This study |
| MGTY096 | TB50a + *pRS426-GFP-2xPH(PLCδ)* + *pRS415-mCh-P4C* | This study |
| JK006 | TB50a *URA3::pRS406-mCh-2xPH(PLCδ)* | This study |
| MTY038 | TB50a *HIS3::pRS403-P(TEF1)-GFP-D4H URA3::pRS406-mCh-2xPH(PLCδ)* | This study |
| MGTY098 | TB50 *lam2Δ::Nat lam4Δ::Nat HIS3::pRS403-P(TEF1)-GFP-D4H URA3::pRS406-mCh-2xPH(PLCδ)* | This study |
| MGTY100 | TB50α *TOR1-1 AVO3-ΔCT::Hph HIS3::pRS403-P(TEF1)-GFP-D4H URA3::pRS406-mCh-2xPH(PLCδ)* | This study |

**Supplementary Table 1:** List of yeast strains used in this study.

| plasmid | source |
| --- | --- |
| pRS426-GFP-2xPH(PLCδ) | PMID: 1185441 |
| pRS414-pTdh3-mCherry-2xPH(PLCδ) | This study |
| pRS416-LactC2-GFP | PMID: 18187657 |
| pRS415-mCh-P4C | Gift from Christopher Stefan lab |
| pRS406-mCh-2xPH(PLCδ) | This study |
| pRS403-P(TEF1)-GFP-D4H | This study |

**Supplementary Table 2:** List of plasmids used in this study.

| substance | | company | cat. no. | stock |
|---|---|---|---|---|
| Rapamycin | | LC laboratories or Thermo Fisher | R-5000 or J62473.MF | 200 µM in DMSO |
| **SAR screen substance** | **carbon tail** | **company** | **cat. no.** | **stock** |
| Carnitine | - | Sigma | C0283 | 10 mM in PBS |
| Palmitate | C16 | Sigma | P0500 | 10 mM in DMSO |
| Palmitoylglycine | C16 | Sigma | 870817P | 10 mM in DMSO |
| Palmitoylglycerol | C16 | Sigma | 75614 | 10 mM in DMSO |
| Palmitoylcholine | C16 | Cayman | Cay9003456-5 | 10 mM in DMSO |
| PAF | C16 | Enzo Life Sciences | BML-L100-0005 | 10 mM in DMSO |
| LauroylCarnitine | C12 | Sigma | 39953 | 10 mM in MeOH |
| MyristoylCarnitine | C14 | Sigma | 61367 | 10 mM in MeOH |
| PalmitoylCarnitine | C16 | Sigma | P4509 | 10 mM in DMSO, or MeOH (only SAR screen) |
| StearoylCarnitine | C18 | Sigma | 61229 | 10 mM in MeOH |
| OleoylCarnitine | C18:1 | Sigma | 870852P | 10 mM in MeOH |
| L-*erythro*-DHS | C16 | Brunschwig | CAY24374 | 5 mM in DMSO |
| D-*erythro*-DHS | C16 | Brunschwig | CAY10007945 | 5 mM in DMSO |
| PHS | C16 | Sigma | P2795 | 10 mM in DMSO |

**Supplementary Table 3:** List of treatment substances and drugs used in this study.

| epitope | type | species | dilution | reference | producer |
|---|---|---|---|---|---|
| Sch9 (total) | polyclonal | Rabbit | 5'000x | | Agrobio - homemade |
| Sch9 pS758 | monoclonal | Mouse | 2'500x | Wanke *et al.*, 2008 | Agrobio - homemade |
| Ypk1 (total) | polyclonal | Rabbit | 25'000x | Bourgoint *et al.*, 2019 | Agrobio - homemade |
| Ypk1 pT662 (1H3) | monoclonal | Mouse | 500x | Berchtold *et al.*, 2012 | IGBMC |
| Akt1 (total) | monoclonal | Mouse | 1'000x | #2920 | Cell Signaling Technology |
| Akt1 pS473 | monoclonal | Rabbit | 750x | #4060 | Cell Signaling Technology |
| anti-Rabbit (H+L) | IRDye 680 (red) conjugated | Donkey | 10'000x | 926-68073 | Li-Cor Biosciences |
| anti-Rabbit (H+L) | IRDye 800 (green) conjugated | Donkey | 10'000x | 926-32213 | Li-Cor Biosciences |
| anti-Mouse (H+L) | IRDye 680 (red) conjugated | Donkey | 10'000x | 926-68072 | Li-Cor Biosciences |
| anti-Mouse (H+L) | IRDye 800 (green) conjugated | Donkey | 10'000x | 926-32212 | Li-Cor Biosciences |

**Supplementary Table 4:** List of antibodies used in this study.
